# Site-Specific Introduction of Alanines for the NMR Investigation of Low-Complexity Regions and Large Biomolecular Assemblies

**DOI:** 10.1101/2023.05.08.539737

**Authors:** Carlos A. Elena-Real, Annika Urbanek, Lionel Imbert, Anna Morató, Aurélie Fournet, Frédéric Allemand, Nathalie Sibille, Jerome Boisbouvier, Pau Bernadó

**Affiliations:** Centre de Biologie Structurale (CBS), Université de Montpellier, INSERM, CNRS. 29 rue de Navacelles, 34090 Montpellier (France); Univ. Grenoble Alpes, CNRS, CEA, Institut de Biologie Structurale (IBS), 71, avenue des martyrs, F-38044, Grenoble, France

**Author notes:** **Corresponding Authors:** Pau Bernadó, Centre de Biologie Structurale (CBS). and Jerome Boisbouvier, Institut de Biologie Structurale (IBS). Equal Contribution (Carlos A. Elena-Real, Annika Urbanek, Lionel Imbert).

**Keywords:** alanine, poly-alanine, isotopic labeling, Nuclear Magnetic Resonance (NMR), methyl-NMR

## Abstract

NMR studies of large biomolecular machines and highly repetitive proteins remain challenging due to the difficulty of assigning signals to individual nuclei. Here, we present an efficient strategy to address this challenge by engineering a *Pyrococcus horikoshii* tRNA/alanyl-tRNA synthetase pair that enables the incorporation of up to three isotopically labeled alanine residues in a site-specific manner using *in vitro* protein expression. We have demonstrated the general applicability of this approach for NMR assignment by introducing isotopically labeled alanines into four proteins, including the 300-kDa molecular chaperone ClpP and the alanine-rich Phox2B transcription factor. For large protein assemblies, our labeling approach enables unambiguous assignments, while avoiding potential artefacts induced by site-specific mutations. When applied to Phox2B, which contains two poly-alanine tracts of nine and twenty alanines, we observe that the helical stability is strongly dependent on the homorepeat length, demonstrating structural cooperativity. The capacity to selectively introduce alanines with distinct labeling patterns is a powerful tool to probe structure and dynamics of biomolecular systems that are out of the reach of traditional structural biology methods.

## INTRODUCTION

Nuclear Magnetic Resonance (NMR) has become a fundamental tool for the high-resolution structural, dynamic and functional characterization of biomolecules in solution. Indeed, NMR is nowadays routinely applied for the investigation of all types of biomolecules from very large biomolecular assemblies (up to 1 MDa) to intrinsically disordered proteins (IDPs) and nucleic acids^[1– 4]^. However, the power of NMR to reveal this detailed information relies on the capacity to identify resonance frequencies of individual nuclei. In this context, the availability of high magnetic fields in combination with multidimensional pulse sequences and isotopic labeling strategies have reduced the signal overlap of the resulting spectra, facilitating the subsequent frequency assignment^[5–7]^. Well-established procedures have been designed for the unambiguous frequency assignment of backbone nuclei of small and medium size proteins (up to ≈80 kDa). All these strategies are based on the specific spectroscopic features of natural amino acids and their connectivity, which emerge through couples of multidimensional heteronuclear experiments^[8]^. Therefore, the absence of signal for certain nuclei, due for instance to chemical exchange processes, or the degeneracy of two or more frequencies can hamper the assignment process. A clear example of severe signal overlap are low-complexity regions (LCRs) in proteins, which present a strong compositional bias that reduces the diversity of chemical environments and, as a consequence, the chemical shift dispersion^[9]^. The site-specific NMR frequency assignment of large biomolecular assemblies is also extremely challenging. Due to their slow tumbling in solution, most of the backbone signals broaden beyond detection, even when using high temperatures and/or perdeuteration. The structural and dynamic investigation of these macromolecular machines has been accomplished by using methyl-NMR^[10–14]^. The fast rotation of the methyl group and the presence of three equivalent hydrogen atoms makes methyl-NMR experiments especially suited to study large biomolecular assemblies. In order to obtain appropriate samples for these experiments, one or a few methyl-containing amino acids (leucine, isoleucine, valine, alanine, threonine and methionine) are incorporated in their ^13^CH_3_-labeled form, while the rest of the protein is deuterated^[15– 17]^. This labeling scheme can be achieved by supplying the bacterial culture with the appropriate isotopologues or with molecules that are precursors in the enzymatic chain of the amino acid synthesis^[18–21]^. However, in methyl-labeled samples, traditional sequential assignment strategies are not possible. For these large machines, mutagenesis-based^[22,23]^ or “*divide-and-conquer*”^[14,24]^ approaches are normally applied. These are costly and laborious approaches and prone to errors caused by local structural perturbations induced by mutations or changes in protein constructs^[22,23]^.

In order to overcome the above-mentioned limitations, we have developed the site-specific incorporation of up to three isotopically labeled alanines into proteins to be applied in large macromolecular machines and LCRs. To achieve this aim, we combined tRNA nonsense suppression with cell-free (CF) protein synthesis^[25]^. We show that the application of this strategy enormously reduces the complexity of the recorded spectra, enabling the study of biomolecular systems not amenable to traditional NMR approaches. Concretely, we have demonstrated our approach for the tetradecameric ClpP protease and the disordered alanine-rich C-terminal region of the Phox2B transcription factor. These proteins represent prototypical examples of very large macromolecular machines and highly degenerated sequences, respectively. On the one hand, ClpP protomers self-assemble into two stacked heptameric rings to form a 300 kDa barrel-shaped serine protease containing a single degradation chamber involving all fourteen copies of the catalytic triad. These proteases play an important role in protein homeostasis and are conserved in bacteria, chloroplasts and mitochondria^[26,27]^. *In vivo*, ClpP forms complexes with ClpX or ClpA, AAA+ ATPases that translocate client proteins through axial pores in the ClpP degradation chamber. On the other hand, the C-terminal region of Phox2B is an example of a LCR rich in alanines, glycines, serines and prolines. Indeed, it contains two poly-alanine (poly-A) tracts of 9 and 20 alanines, which is the longest one found in the human proteome. Interestingly, the expansion of the long poly-A tract with +5, +7 or +11 alanines causes a severe developmental pathology, the Congenital Central Hypoventilation Syndrome (CCHS)^[28,29]^. The structural changes occurring in Phox2B upon the alanine expansion that trigger the pathology remain unknown, equivalently to eight other rare diseases linked to the abnormal expansion of poly-A tracts^[30,31]\^

## RESULTS AND DISCUSSION

The site specific isotopic labeling (SSIL) approach that we have developed uses a synthetic nonsense suppressor tRNA (tRNA_CUA_) and aminoacyl-tRNA synthetase (aaRS) pair that should be orthogonal to the *Escherichia coli* translation system^[25]^. tRNA_CUA_ enables the insertion of a labeled amino-acid specifically at the position encoded by an amber stop codon, while its orthogonality to endogenous tRNA/aaRS pairs prevents its reloading with the non-labeled amino acid present in the CF reaction. To incorporate alanines, a previously characterized tRNA/alanyl-RS (AlaRS) pair from *Pyrococcus horikoshii* was chosen^[32,33]^ with the anticodon mutated to obtain the tRNA_CUA_. First, the capacity of the recombinant AlaRS to load the synthetic tRNA_CUA_ with alanine was evaluated using PAGE. When optimal conditions were used, an aminoacylation level of ∼50% was achieved (see Fig. S1 in Supplementary Information). The orthogonality of this pair was subsequently tested by monitoring the CF expression of huntingtin exon-1 protein containing 16 consecutive glutamines fused to superfolder green fluorescent protein (HttExon1) in which the A53 codon was exchanged by the amber stop codon (HttExon1-A53). When the tRNA_CUA_ was used in the absence of the AlaRS, the protein was synthesized at similar levels as in the positive control, without the amber stop codon (Fig. 1A). This observation indicated that the tRNA_CUA_ could be loaded by the endogenous *E. coli* AlaRS, showing that the *P. horikoshii* tRNA_CUA_ was not orthogonal to the *E. coli* translational machinery. To improve the required orthogonality, we engineered the *P. horikoshii* pair. Previous studies highlighted the relevance of the highly conserved G3:U70 base pair located in the acceptor stem of the tRNA^Ala^ for the selective aminoacylation with alanine^[34,35]^. In a recent study, it was shown that the substitution of this base pair by G3:C70 impaired alanine loading by *E. coli* AlaRS^[36]^. Interestingly, the efficient aminoacylation of the G3:C70 tRNA^Ala^ was recovered when highly conserved Asn303 and Asp400 residues in *E. coli* AlaRS were mutated to alanine. Based on these observations, we designed a new *P. horikoshii* tRNA_CUA_/AlaRS pair with G3:C70, which would hamper the recognition by *E. coli* AlaRS, and two alanines at positions corresponding to Asn360 and Asp459 in the *P. horikoshii* AlaRS, enabling an efficient external loading of the tRNA_CUA_. Using similar aminoacylation conditions as for the wild-type pair (see methods section), ∼50% of loaded tRNA_CUA_ was obtained, suggesting a minimal impact of the designed mutations in the *P. horikoshii* pair (Fig. 1B). Importantly, in the absence of the mutant AlaRS, a substantial reduction of HttExon1-A53 CF expression was observed (Fig. 1A), indicating an improvement in the tRNA_CUA_ orthogonality upon introducing the G3:C70 base pair. Addition of the mutant AlaRS to the CF reaction increased the protein expression, evidencing the recognition of suppressor tRNA_CUA_ by the enzyme under the experimental conditions, which notably differ from those found for the optimal external loading. When increasing amounts of aminoacylated tRNA_CUA_ were added to the CF reaction, protein synthesis improved, reaching its maximum at 20 μM of loaded tRNA_CUA_ (Fig. 1C). As previously observed^[25]^, further increasing the tRNA_CUA_ concentration compromised the protein synthesis. In order to validate our approach for NMR studies, a 5 mL CF reaction was prepared using 10 μM of [^13^C,^15^N]-alanine loaded tRNA_CUA_ to express HttExon1-A53. After purification, the ^15^N-HSQC spectrum showed a single correlation that overlapped with that previously assigned for A53 using multidimensional NMR (Fig. 1D)^[37]^.

**Figure 1.**
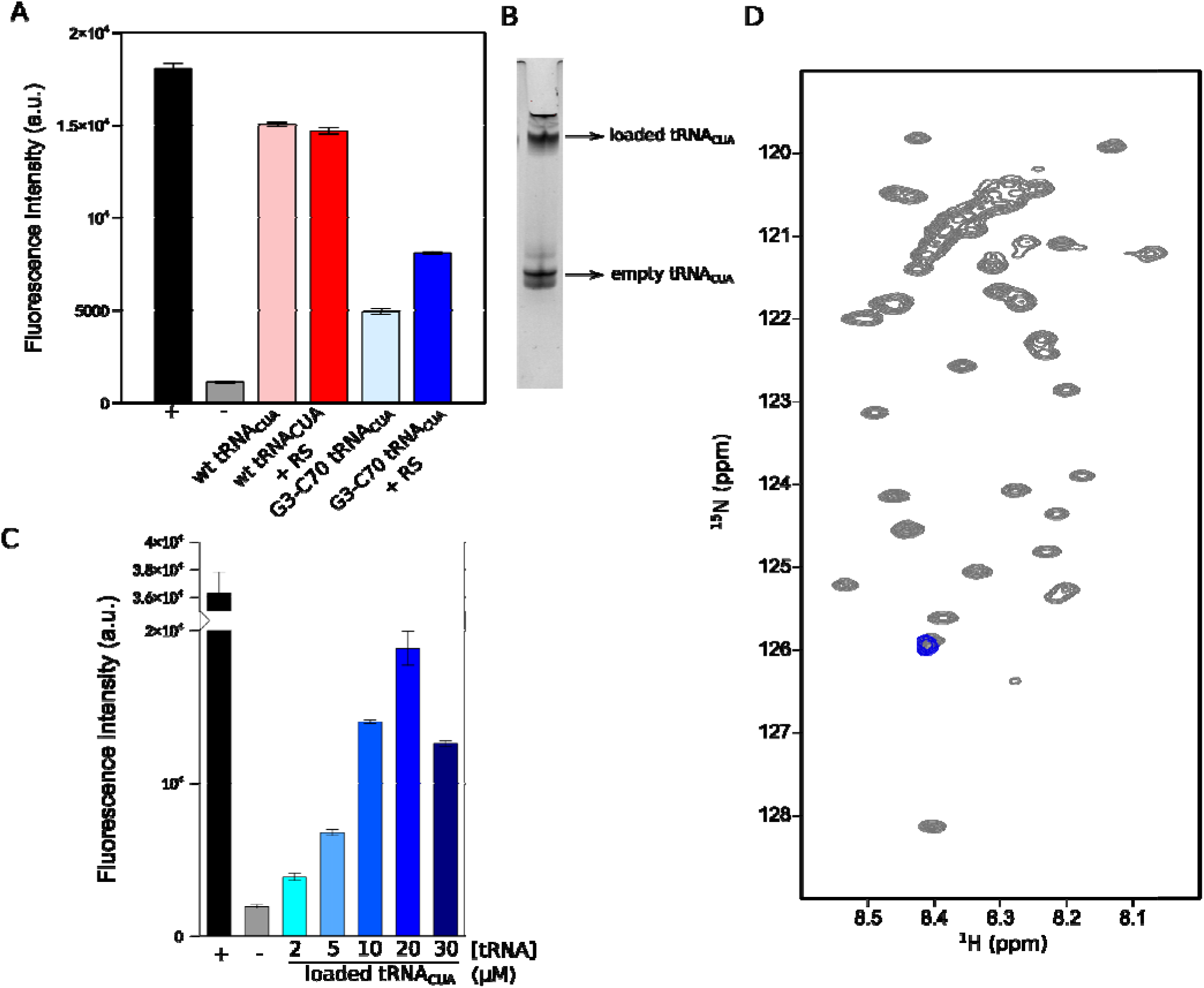
**(A)** Endpoint fluorescence intensity plot for CF expression of HttExon1-A53 using wt or G3:C70 tRNA_CUA_ and in the absence or presence of AlaRS. + and – indicate the positive (wild-type HttExon1) and negative (HttExon1-A53 without tRNA_CUA_) controls. **(B)** SDS-PAGE monitoring the efficiency (∼50%) of the suppressor tRNA_CUA_ loading. **(C)** Titration of CF reactions with increa ing concentrations of alanine-loaded tRNA_CUA_. **(D)** ^15^N-HSQC spectrum of HttExon1 fully labeled (grey) and HttExon1-A53 SSIL sample (blue) where a single peak, corresponding to A53 was observed.

The ability to produce isotopically labeled samples in a site-specific manner allows the quantification of the orthogonality improvement when using the mutated *P. horikoshii* tRNA_CUA_/AlaRS couple. For this, two samples of the nucleotide-binding domain (NBD) of the HMA8 ATPase^[38]^ were prepared using the aminoacylated wild-type or the G3:C70 mutated tRNA_CUA_. A third reference sample was labeled on all the alanine methyl sites. To quantify the level of incorporation of ^13^CH_3_-labeled alanine at the amber codon-encrypted site, the three samples were produced in the presence of ^13^CH_3_-labeled methionine to serv1e as an internal reference. Comparison of signal intensities with the reference sample revealed that the incorporation level of ^13^CH_3_-alanine encoded by the amber codons was increased by a factor 3.2 when the G3:C70 mutation was introduced in the tRNA_CUA_ (Fig. S2). Although the incorporation was not complete (60.2%), it enabled the systematic application for NMR assignment.

The above-described results validate the use of the engineered *P. horikoshii* tRNA_CUA_/AlaRS pair as a tool to isotopically label single alanines in a site-specific manner. Next, we applied this technology to study ClpP. First, we analyzed the signal dispersion of alanines in this macromolecular complex. For this, ClpP was produced *in vitro* in D_2_O-based buffer and in the presence of a mixture of perdeuterated amino acids, supplemented with an excess of [2-^2^H,3-^13^C]-L-alanine. This procedure yielded an optimally labeled sample for the acquisition of high-resolution 2D-methyl TROSY experiments (Fig. 2A)^[39,40]^. Although most of the 24 alanine methyl signals were well resolved, the characteristically slow overall tumbling of this 300 kDa particle precluded the acquisition of triple-resonance experiments necessary for sequence-specific assignment. To test the SSIL strategy in the context of a large protein assembly, we targeted two alanines with expectedly different features. On the one hand, A96 is located in a globular section of the complex, in close proximity of the active site. On the other hand, a more flexible behavior was expected for A11, which is involved in ClpP gating and target recognition^[10]^. To identify these two residues, we used the above-described nonsense suppressor tRNA_CUA_ loaded with [2-^2^H,3-^13^C]-L-alanine to prepare two ClpP samples. A well-resolved alanine signal was observed in both cases (Fig. 2B,C), allowing direct assignment of the two methyl probes. The low intensity observed for the A11 methyl signal is presumably due to an exchange between different conformations of the N-terminal gate segment of ClpP^[41]^. Note that this approach does not suffer from the secondary chemical shift perturbation as observed using mutagenesis-based approaches^[22]^.

**Figure 2.**
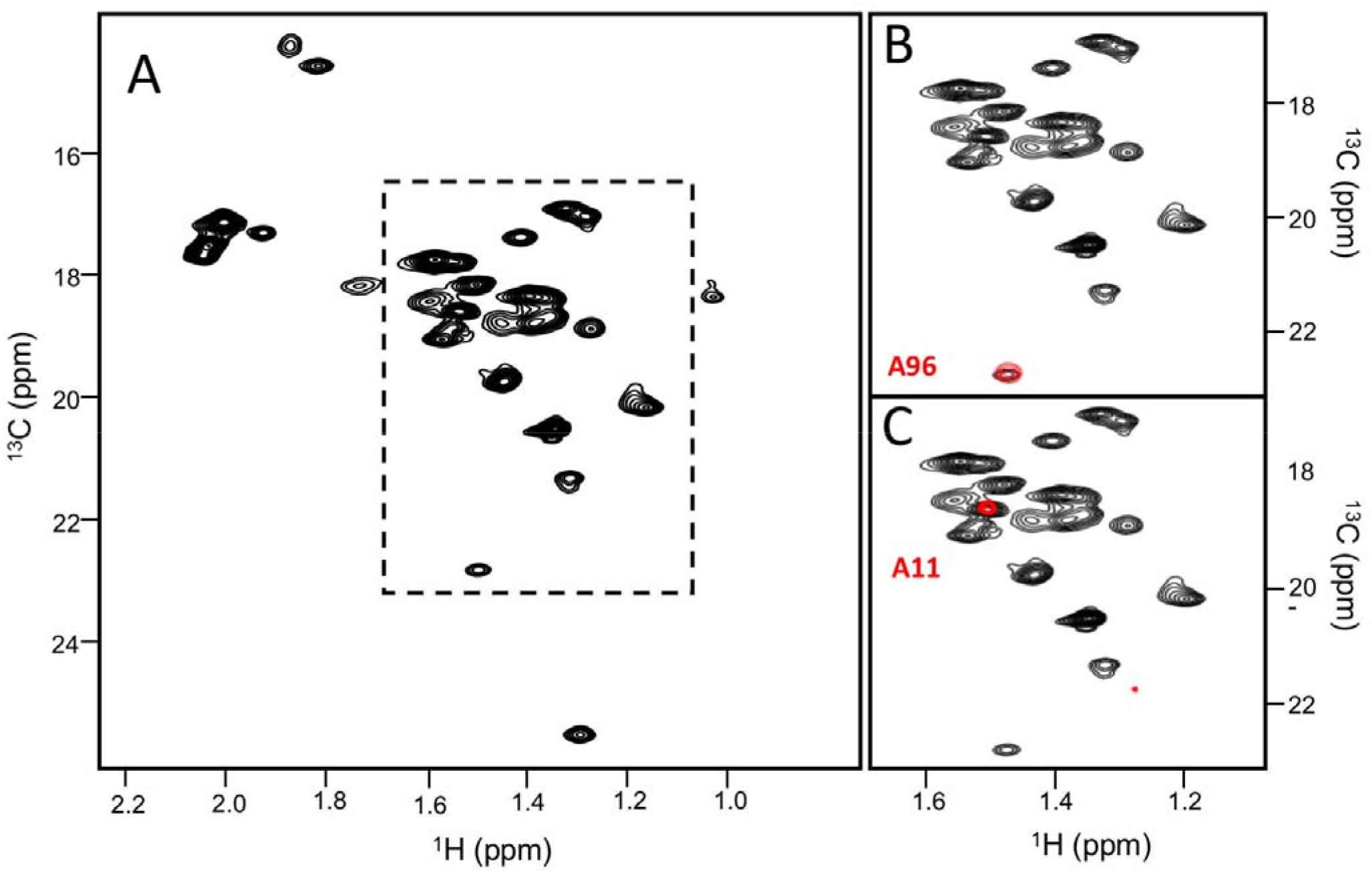
**2D methyl TROSY of the 300 kDa ClpP. (A)** Reference spectrum of perdeuterated ClpP where all alanines and methionines were ^13^CH_3_-labeled. Zoom corresponding to the dashed delimited reference spectrum overlayed with SSIL spectra (red) where a single alanine methyl signal was observed for A96 **(B)** and A11 **(C)**

Next, we applied the alanine SSIL strategy to investigate the two poly-A tracts present at the C-terminal domain of Phox2B. This 179-residue long fragment (from residue 136 to 314) is a LCR highly enriched in alanine, glycine, serine and proline, which yielded a severely overlapped ^15^N-HSQC spectrum (Fig. S3). When the protein was produced in CF using [^13^C,^15^N]-L-alanine as the only isotopically enriched amino acid, the ^15^N-HSQC exhibited multiple alanine signals, with many of them clustering in a narrow region of the spectrum (Fig. S3A). Similarly, the Cα-Hα and Cβ-Hβ regions of the ^13^C-HSQC displayed few connected broad signals and no isolated peaks could be identified (Fig. S3). The ensemble of these observations indicated that traditional frequency assignment with multidimensional NMR spectra was not applicable to characterize this protein.

In order to overcome the severe overlap, we applied our strategy to study two alanines located at the center of the short (A163) and long (A250) poly-A tracts of Phox2B (Fig. 3A). Two NMR samples were prepared by introducing a [^13^C,^15^N]-L-alanine in these two positions, while keeping all the other amino acids unlabeled. Note that deuterated alanine was used in the CF reaction in order to avoid spurious signals arising from natural abundance^[42]^. Well-resolved signals were obtained for N-H, Cα-Hα and Cβ-Hβ correlations (Fig. 3B,C and D), demonstrating the power of SSIL to resolve degenerated frequencies in homorepeats. Interestingly, despite the fact that both alanines, A163 and A250, experience the same sequence context, they displayed different chemical shifts for the three studied correlations. Indeed, the largest differences between both alanines were observed for the Cα and Cβ frequencies, which are the most sensitive atoms to secondary structure propensities. To further demonstrate this point, a secondary chemical shift (SCS) analysis was performed by comparing the experimental Cα and Cβ chemical shifts with these derived from a neighbor-corrected random coil library^[43]^. The SCS analysis revealed that both A163 and A250 adopt a helical conformation, although with a different propensity, with A250 exhibiting a higher structuration level (3.3 ppm) than A163 (1.5 ppm).

**Figure 3.**
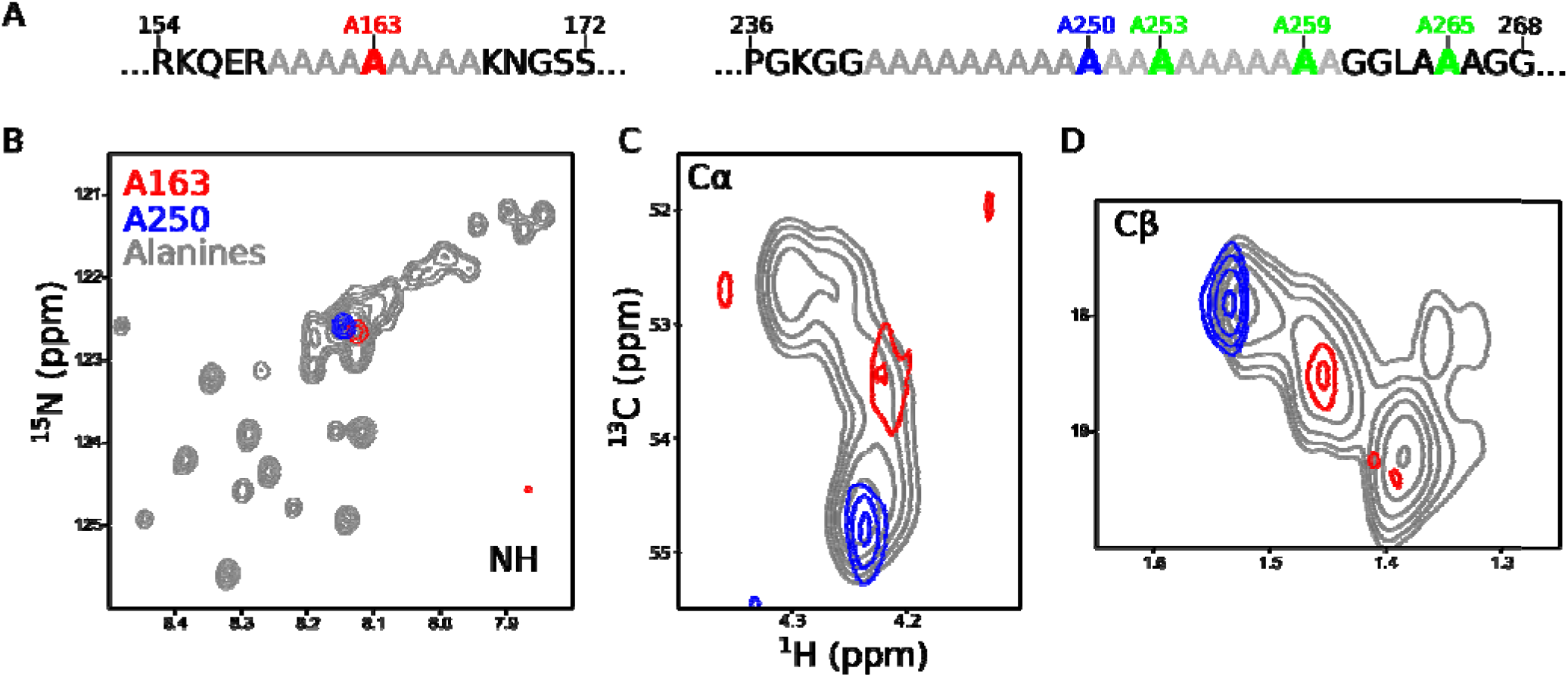
(**A)** Sequences of short (left) and long (right) poly-A stretches present in Phox2B C-terminal domain. Alanines probed in this study are highlighted. Overlays of the single alanine ^15^N-HSQC and ^13^C-HSQC spectra of Phox2B C-terminal domain measured for samples with all alanines ^15^N,^13^C-labeled (grey), showing NH **(B)**, Cα-Hα **(C)** and Cβ-Hβ **(D)** correlations. Red (A163) and blue (A250) colors are consistent in all panels.

In order to further explore the effect of the poly-A length on the helical stability, we studied three additional alanines: A253 and A259, which are located in the long poly-A tract, and A265, which is placed in the center of a short three-alanine segment next to the long tract (Fig. 3A). For this assignment we used the recently described multi-SSIL approach^[44]^, by simultaneously incorporating three isotopically labeled alanines into Phox2B. In order to increase the protein yield and obtain a 7 μM sample, 20 μM of loaded tRNA_CUA_ was added to 10 mL of CF reaction. Three isolated frequencies could be identified in the NH, Cα-Hα and Cβ-Hβ regions of the spectra, overcoming the severe overlap observed when all alanines of Phox2B were isotopically labeled. In order to assign these correlations, we produced two other multi-SSIL samples in which only one of the three labeled alanines coincided with the original one. Concretely, A242, *A253* and A264 were isotopically labeled in the first sample, while A159, *A259* and A266 were labeled in the second one (Fig. 4A). This strategy enabled us to unambiguously assign the three targeted alanines. Each of the ^15^N-HSQC spectra of these two additional samples yielded three isolated NH frequencies and only one of them overlapped with the original spectrum, enabling the direct assignment of A253 and A259, while the only unconnected frequency was identified as A265. Due to the limited carbon frequency dispersion in Phox2B, a slightly different scenario occurred for Cα-Hα and Cβ-Hβ for which only two isolated peaks were observed in some of the spectra. Despite the reduced number of isolated frequencies, the overlap of concatenated spectra also yielded the unambiguous assignment of Cα-Hα and Cβ-Hβ correlations for A253, A259 and A265. Note, however, that further concatenation could produce some assignment ambiguities, which could be eventually overcome with strategically designed singly labeled alanine samples. Therefore, despite the very limited frequency dispersion of alanine carbon correlations in Phox2B, these experiments demonstrate that the concatenation of multi-SSIL samples is feasible with the engineered *P. horikoshii* tRNA_CUA_/AlaRS pair.

**Figure 4.**
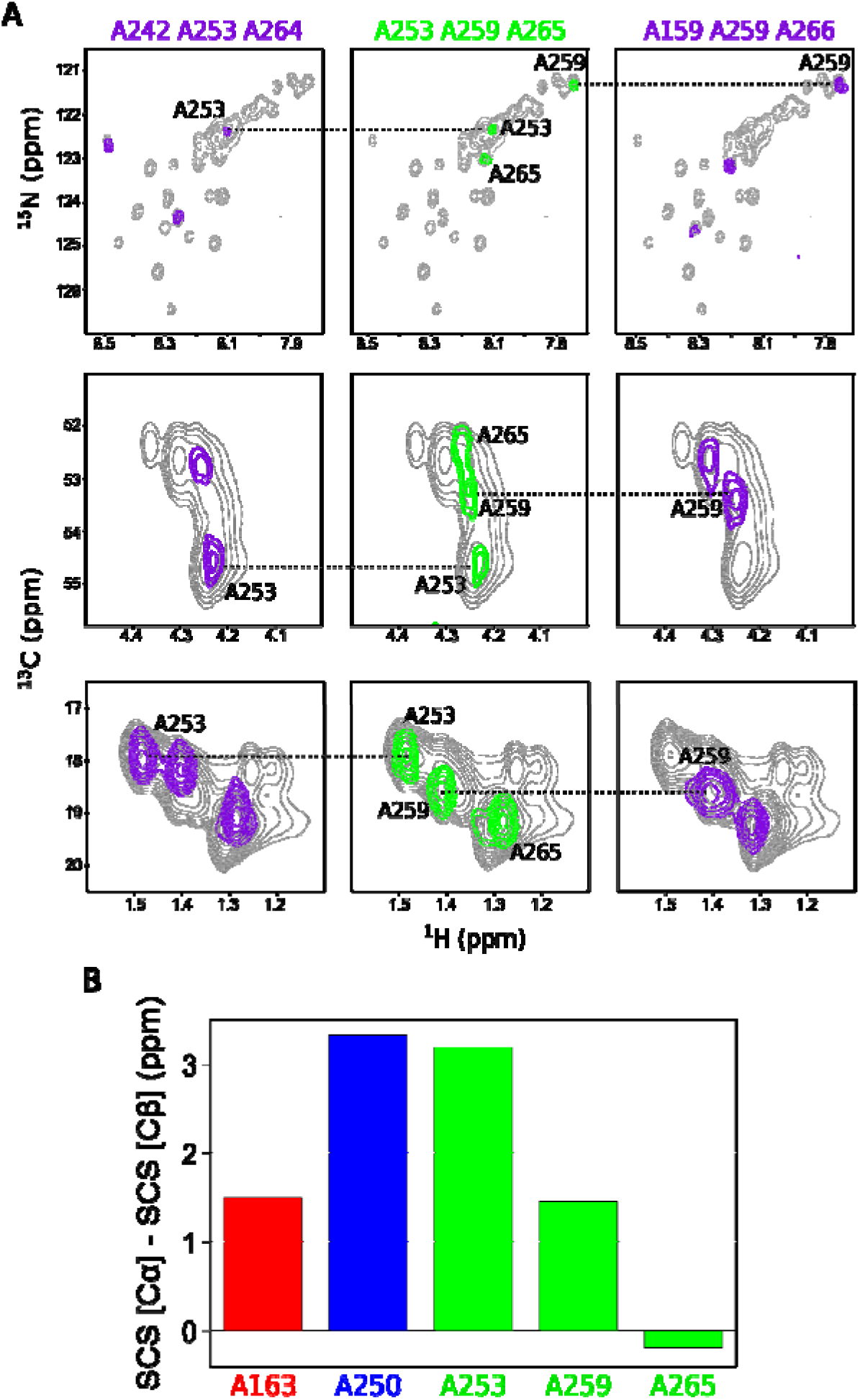
**(A)** Zooms of the NMR spectra displaying the NH, Cα-Hα and Cβ-Hβ correlations (from top to down) for the A253-A259-A265 specifically labeled sample (in green) and its two concatenated samples (A242-*A253*-A264, purple, left) and A159-*A259*-A266 (purple, right) that enable the unambiguous assignment. Horizontal dashed lines indicate the common peaks in two concatenated spectra. **(B)** Secondary chemical shift (SCS) analysis using experimental Cα and Cβ chemical shifts and a random-coil library. A163 (red), and A250 (blue) measured using SSIL, and A253, A259 and A265 (green) measured in a single sample using multi-SSIL

We then performed an SCS analysis including the three additional alanines, which displayed very different helical propensities (Fig. 4B). Similarly to A250, A253, which is also located in the central region of the long poly-A tract, displayed a strong helical propensity. Conversely, A259, which is positioned towards the C-terminus of the same tract, exhibited a reduced helical behavior. Indeed, this terminal alanine had a structuration level very similar to that observed for A163, located in the center of the short poly-A tract. Interestingly, A265, which probed the helical propensity of the three-alanine stretch, displayed a random coil conformational behavior.

Next, we investigated the helical fraction (HF) of the poly-A tracts of Phox2B. The precise calculation of the HF from sparse chemical shift data is difficult. Cα chemical shifts for several poly-A tracts in a forced α-helical conformation thanks to designed N- and C-capping residues have been experimentally determined, indicating a deviation of 2.3 ppm from random coil estimated values^[45]^. Assuming a linear relationship between chemical shift deviation and helicity^[46]^, our Cα data indicate that the long poly-A tract of Phox2B has an HF of ∼90% at the central region that decreases to ∼25% at its C-terminus (A259). Interestingly, the short poly-A tract displays a maximum HF of ∼25%.

This structural propensity found in our NMR investigation of Phox2B is in line with the pioneering study of Gratzer and Doty^[47]^ and two recent NMR studies that unambiguously identified three stretches with five, six and eight consecutive alanines as partially α-helical^[48,49]^. The ensemble of these studies on poly-A in their protein context indicates a positive correlation between homorepeat length and helical propensity. These observations are in contradiction with other studies on synthetic poly-A peptides in which the residues were identified as disordered with some prevalence for extended poly-proline-II conformations^[50,51]^. Furthermore, our study demonstrates a cooperative effect stabilizing this secondary structure when incorporating additional alanines in the homorepeat. This increased helical length and stability could be key in enhancing protein self-interaction through coiled-coils^[52]^ and trigger aggregation^[53]^ or phase separation^[54]^ of pathogenic forms of poly-A-hosting proteins. In addition to the inherent α-helical propensity of poly-A tracts, it has been recently suggested that, in a protein context, this conformational behavior could be tuned by the nature of the flanking residues, where the presence of N- or C-capping residues could further stabilize the inherent helical propensity of the poly-A tracts^[55]^.

In summary, despite the technical advances in multiple fronts, the high-resolution investigation of large biomolecular machines and LCRs by NMR remains a challenge. Here, we have presented an engineered tRNA_CUA_/AlaRS pair enabling the site-specific incorporation of up to three alanines in four distinct proteins in terms of size, structure and sequence composition, demonstrating the generality of the approach. With this methodology, the assignment of biological machines is facilitated and rendered independent of the specific mutation introduced in mutagenesis-based approaches. Furthermore, it enables the structural and dynamic characterization of alanine-rich proteins, including transcription factors causing poly-A-expansion developmental and neurodegenerative diseases. The extension of the SSIL strategy to other amino acids will permit structural and functional studies of other challenging relevant biomolecular systems, including the kinetic investigation of large catalytic machines or the structural bases of the emerging phenomenon of LCR-driven liquid-liquid phase separation^[56,57]^.

## Supporting information

Experimental Protocols and supplemental figures S1-3

## Acknowledgements

This work was supported by the European Research Council under the European Union’s H2020 Framework Programme (2014-2020)/ERC Grant agreement n° [648030], and MUSE-App 2021 Ondine ANR-16-IDEX-0006 awarded to PB and ProteaseInAction ANR-19-CE11-0022 awarded to JB. The CBS is a member of France-BioImaging (FBI) and both CBS and IBS are members of the French Infrastructure for Integrated Structural Biology (FRISBI), supported by ANR-10-INBS-04-01 and ANR-10-INBS-05 grants, respectively. This work used facilities at the Grenoble Instruct-ERIC Center (ISBG; UAR 3518 CNRS-CEA-UGA-EMBL) within the Grenoble Partnership for Structural Biology (PSB), with support of GRAL, a project of the University Grenoble Alpes graduate school (Ecoles Universitaires de Recherche) CBH-EUR-GS (ANR-17-EURE-0003). IBS acknowledges integration into the Interdisciplinary Research Institute of Grenoble (IRIG, CEA).

