## Supplementary material for "Site-Specific Introduction of Alanines for the NMR Investigation of Low-Complexity Regions and Large Biomolecular Assemblies": Experimental Protocols and supplemental figures S1-3

### Equal Contribution

#### Experimental Protocols and Supplemental Figures

Unless specified otherwise, all chemicals were obtained from Sigma-Aldrich (St. Quentin Fallavier, France).

##### Protein constructs for suppression samples

All plasmids were prepared as previously described[1]. Synthetic genes of wild-type huntingtin exon1 with 16 consecutive glutamines (HttExon1Q16), Phox2B C-terminal domain (residues from 136 to 314) and carrying amber codons (TAG) instead of alanines codons e.g. A53 (HttExon1Q16-A53) were ordered from GeneArt® (ThermoFisher Scientific, Illkirch, France). All genes were cloned into pIVEX 2.3d 3C-sfGFP-$His_6$. Synthetic genes of ClpP from *Thermus thermophilus*, wild-type and mutants carrying amber codons (TAG) instead of alanine codons, *e.g.* A96-ClpP or A11-ClpP, were ordered from Genecust (Boynes, France). The sequence of all plasmids was confirmed by sequencing by GENEWIZ® (Leipzig, Germany).

##### Standard cell-free expression conditions

Lysate was prepared as previously described[1]. It is based on the *Escherichia coli* strain BL21 Star (DE3)::RF1-$CBD_3$, generously provided by Prof. Gottfried Otting (Australian National University, Canberra, Australia)[2]. Cell-free protein expression was performed as described in [3,4]. Nevertheless, the magnesium acetate (5-20 mM) and potassium glutamate (60-200 mM) concentrations were optimized for each new batch of S30 extract. A titration of both compounds was performed to obtain the maximum yield.

##### Preparation of aminoacylated suppressor $tRNA_{CUA}$

A $tRNA_{CUA}$/tRNA synthetase pair from *Pyrococcus horikoshii*[5,6] was prepared in house. Substitution of the G3:U70 by a G3:C70 base pair in the $tRNA_{CUA,}$ as well as N360A and E459A mutations on the alanine tRNA synthetase (AlaRS) were designed to improve orthogonality[7]. The optimized genes encoding the wild-type and double mutation (N306A and E459A) of truncated N752 *P. horikoshii* Alanyl-tRNA synthetase (*Ph* AlaRS) were ordered from IDT DNA Technologies. These genes were cloned in pET26b vector between *NdeI* and *XhoI* sites in frame with the $(HIS)_6$-tag at the C-terminal end. Plasmids were transformed into *E. coli* BL21(DE3) strain. Wild-type and mutated proteins were expressed in *E. coli* BL21(DE3) cells in LB medium at 37°C supplemented with 50 µg/mL Kanamycin. Protein expression was induced by IPTG (0.5 mM) and cells were grown for 3 hours before harvesting them by centrifugation for 20 min at 6000 xg at 4°C. The pellet was resuspended in 20 mM Tris-HCl pH 7.5, 300 mM NaCl and 2 mM DTT (buffer A) and stored at -80°C. Cells were supplemented with protease inhibitors (cOmplete™ EDTA-free protease inhibitor cocktail), lysed by sonication, and insoluble proteins and cell debris were sedimented by centrifugation at 40000 xg at 4°C for 30 min. The soluble fraction was aliquoted in 2 mL tubes, heated at 70°C for 20 min and centrifuged at 15000 xg at RT for 10 min. The supernatant was pooled and supplemented with imidazole to a final concentration of 10 mM, filtered through 0.45 µm filters and loaded onto an affinity column (5 mL HisTrap™ Excel, Cytiva), equilibrated with buffer B (buffer A containing 10 mM imidazole). The column was washed with buffer B and proteins were eluted with a linear 0–100% gradient of buffer C (buffer A containing 0.5 M imidazole). The peak fractions were analysed by SDS-PAGE. Fractions containing tagged *Ph* AlaRS were pooled and dialysed overnight at 4°C against 25 mM Tris-HCl pH 7.5, 5 mM DTT and 50 µM zinc acetate and concentrated to 10 mg/mL.

The suppressor $tRNA_{CUA}$ and the G3:C70 mutant were transcribed *in vitro* and purified by phenol-chloroform extraction. Prior to use, suppressor $tRNA_{CUA}$ was refolded in 100 mM HEPES-KOH pH 7.5, 10 mM KCl at 70°C for 5 min and a final concentration of 5 mM $MgCl_2$ was added just before the reaction was placed on ice. The refolded $tRNA_{CUA}$ was then aminoacylated with [$^{15}$N,$^{13}$C]- L-alanine (CortecNet, Les Ulis, France) or [2-$^{2}$H,3-$^{13}$C] -L-alanine (NMR-Bio, France) in a standard aminoacylation reaction: 20 µM $tRNA_{CUA}$, 4 µM AlaRS (wild-type or mutant), 0.4 mM of the specific alanine in 100 mM sodium acetate pH 5.0, 10 mM KCl, 20 mM $MgCl_2$, 0.5 mM TCEP, 5 mM ATP. After incubation at 45°C for 1 hour, loaded suppressor $tRNA_{CUA}$ was precipitated with 300 mM sodium acetate pH 5.2 and 2.5 volumes of 96% EtOH at -80°C and stored as dry pellets at -20°C. Successful loading was confirmed by polyacrylamide gels with 10% acrylamide (19:1), 6 M urea and 37 mM PIPES pH 6.0. To enable the analysis of empty and loaded tRNA, the free amine of the loaded amino acid was modified with sNHS-biotin and conjugated to streptavidin as reported by Pütz *et al.*[8] and the Suga lab[9,10].

**Optimization of cell-free suppression conditions**

In order to optimize the loaded $tRNA_{CUA}$ concentrations for the CF reaction, a titration was performed with final $tRNA_{CUA}$ concentrations between 0 and 30 µM. Protein expression was monitored by measuring sfGFP fluorescence in a plate reader/incubator (Gen5 v3.03.14, BioTek Instruments, Colmar, France) at 485 nm (excitation) and 528 nm (emission). Assays were carried out in a reaction volume of 50 µL dispensed in 96-well plates and incubated at 23°C for 5 h. A similar test was performed for ClpP.

#### Preparation of NMR samples

Samples for NMR studies were produced by CF at 5-20 mL scale and incubated at 23°C and 450 rpm in a thermomixer for 4 h or at 27°C at 20 rpm in a hybridization oven (Techne) for 3 h for ClpP. Uniformly labeled NMR samples of HttExon1 and Phox2B were obtained by substituting the standard amino acid mix with 3 mg/mL [$^{15}$N,$^{13}$C]-labeled ISOGRO®[11] (an algal extract lacking four amino acids: Asn, Cys, Gln and Trp) and additionally supplying [$^{15}$N, $^{13}$C]-labeled Asn, Cys and Trp (1 mM each) and 2 mM Gln (CortecNet, Les Ulis, France). The perdeuterated ClpP sample specifically $^{13}CH_3$-labelled on all the alanine and methionine residues, was obtained using a mix of 20 [$^{2}$H,$^{15}$N]-labelled amino acids (3 mg/mL; CIL, USA), supplemented with an excess of [2-$^{2}$H,3-$^{13}$C]-L-alanine, (0.5 mg/mL; NMR-Bio, FR) and L-methionine (2,3,3,4,4-$^{2}H_5$, Methyl-$^{13}CH_3$), (0.8 mg/mL CIL, USA). 15 µM of [$^{15}$N,$^{13}$C]-alanine suppressor $tRNA_{CUA}$ was added to perform single site-specific suppression of HttExon1-A53 and Phox2B samples, while 20 µM of [$^{15}$N,$^{13}$C]-L-Alanine loaded $tRNA_{CUA}$ was added for multiple site-specific suppression. For Phox2B suppression samples [$^{2}$H]-L-Alanine and Serine residues (2 mM) were used to eliminate the presence of natural abundance signals. The site specifically labeled $^{13}CH_3$-A11-ClpP and $^{13}CH_3$-A96-ClpP samples were produced *in vitro* using a mix of 20 [$^{2}$H,$^{15}$N]-labelled amino acids (3 mg/mL; CIL, USA), supplemented with 20 µM of suppressor $tRNA_{CUA}$ loaded with [2-$^{2}$H,3-$^{13}$C]-L-alanine. The Nucleotide-Binding Domain of the P1B-type ATPase HMA8 from *Arabidopsis thaliana* (NBD, 16.7 kDa, 155 amino acids) sample specifically $^{13}CH_3$-labelled on all the alanine and methionine residues, was produced in CF using a mixture of the 20 amino acids at a concentration of 1 mM each. [2-$^{2}$H,3-$^{13}$C]-L-alanine (NMR-Bio, FR) and L-methionine (2,3,3,4,4-$^{2}H_5$, Methyl-$^{13}CH_3$, CIL, USA) were mixed with 18 other amino acids in unlabeled form. The site-specifically labeled $^{13}CH_3$-A85-NBD samples were produced *in vitro* using a mixture containing L-methionine (2,3,3,4,4-$^{2}H_5$, Methyl-$^{13}CH_3$, 1 mM), 19 unlabeled amino acids (1 mM each), supplemented with 20 µM of suppressor $tRNA_{CUA}$ (wild-type or mutated) loaded with [2-$^{2}$H,3-$^{13}$C]-L-alanine.

#### Protein sample purification

The purification of HttExon1 and Phox2B C-terminal domain and their suppression mutants was performed at room temperature. The CF reaction was diluted 10-fold with buffer A (50 mM Tris-HCl pH 7.5, 1 M NaCl) before incubating it 1 h with 1.5 mL of Ni-resin (cOmplete™ His-Tag Purification Resin). The matrix was packed by gravity-flow and washed with increasing concentrations of imidazole (10, 15, 25, 50, 100 and 200 mM). Elution fractions were checked under UV light and fluorescent fractions were pooled, protease inhibitors were added (cOmplete™ EDTA-free protease inhibitor cocktail); 1 mM DTT was added to Phox2B C-terminal domain samples. Then, proteins were dialyzed against NMR buffer (20 mM BisTris-HCl pH 6.5, 150 mM NaCl) at 4°C using SpectraPor 4 MWCO 12-14 kDa dialysis tubing (Fisher Scientific, Illkirch, France). Dialyzed proteins were then concentrated with 10 kDa MWCO Vivaspin centrifugal concentrators (3500 xg, 4°C) (Sartorius, Göttingen, Germany). Protein concentrations were determined by means of fluorescence using a sfGFP calibration curve. Final NMR sample concentrations ranged from 8 to 40 µM. Protein integrity was analyzed by SDS-PAGE.

*T. thermophilus* ClpP samples produced by CF were purified using a heat shock (60°C for 20 minutes) followed by chromatography using a Resource Q resin (GE, USA). Elution fractions containing ClpP were concentrated with 50 kDa MWCO Amicon centrifugal concentrators (3500 xg) (Millipore, FR) in the final NMR buffer (99,8% $D_2O$, 20 mM Tris-HCl pH 7.5, 100 mM NaCl, $MgCl_2$ 5 mM). Final NMR sample concentrations ranged from 20 to 400 µM (ClpP monomer concentration). The ATPase HMA8 NBD samples were purified in a single step using Ni-NTA affinity chromatography and concentrated (20 to 100 µM) in the NMR buffer (99,8% $D_2O$, 20 mM pH 8, 100 mM NaCl).

**NMR experiments and data analysis**

All NMR HttExon1 and Phox2B samples contained final concentrations of 10% $D_2O$ and 0.5 mM 4,4-dimethyl-4-silapentane-1-sulfonic acid (DSS). NMR experiments for HttExon1 and Phox2B were performed at 293 K on a Bruker Avance III spectrometer (Bruker Biospin, Wissembourg, France) operating at a $^1$H frequency of 800 MHz. $^{15}$N-HSQC and $^{13}$C-HSQC were acquired in order to determine amide ($^1H_N$ and $^{15}$N) and aliphatic ($^1H_{aliphatic}$ and $^{13}C_{aliphatic}$) chemical shifts, respectively. Spectrum acquisition parameters were set up depending on the sample concentration. For ClpP samples, 2D $^1$H-$^{13}$C SOFAST methyl-TROSY experiments[12] were recorded at 333 K, on a Bruker Avance III HD spectrometer equipped with a 5 mm cryogenically cooled, pulsed-field-gradient triple-resonance probe operating at a $^1$H frequency of 600 MHz. Extra signals corresponding to protein impurities were greatly reduced by subtracting NMR spectra acquired on a control sample produced *in vitro* using same protocol but in the absence of ClpP plasmid. 2D $^1$H-$^{13}$C SOFAST methyl-TROSY experiments[12] for Nucleotide-Binding Domain of ATPase HMA8 were recorded at 298 K, on a Bruker Avance III HD spectrometer equipped with a 5 mm cryogenically cooled, pulsed-field-gradient triple-resonance probe operating at a $^1$H frequency of 850 MHz.

All spectra were processed with TopSpin v3.5 (Bruker Biospin, Wissembourg, France) and analyzed using CCPN-Analysis software v2.4[13]. Chemical shifts were referenced with respect to the $H_2O$ signal relative to DSS using the $^1$H/X frequency ratio of the zero point according to Markley et *al.*[14]

Random coil chemical shifts for Phox2B were predicted using POTENCI, a pH, temperature and neighbour corrected IDP library (https://st-protein02.chem.au.dk/potenci/)[15]. Secondary chemical shifts (SCS) were obtained by subtracting the predicted value from the experimental one (SCS=$\delta_{exp}$-$\delta_{pred}$).

**FIGURE S1.**

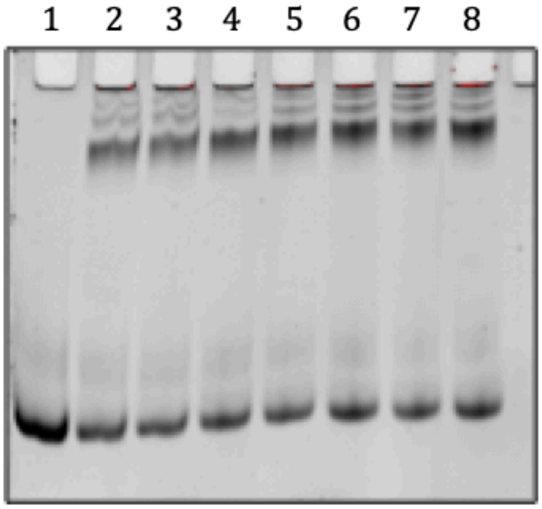


**Figure S1. Time-course alanine loading of non-mutated tRNA$_{CUA}$.** Line (1) corresponds to the empty tRNA$_{CUA}$, subsequent lines correspond to the loading state after 5, 10, 30, 45, 60, 90 and 120 minutes of reaction. Bottom band corresponds to the unloaded tRNA$_{CUA}$ and the top one to the loaded one after modification with sNHS-biotin and conjugated to streptavidin. Reaction performed at pH 7.5 and 37ºC using 2 µM of wild-type *P. horikoshii* AlaRS.

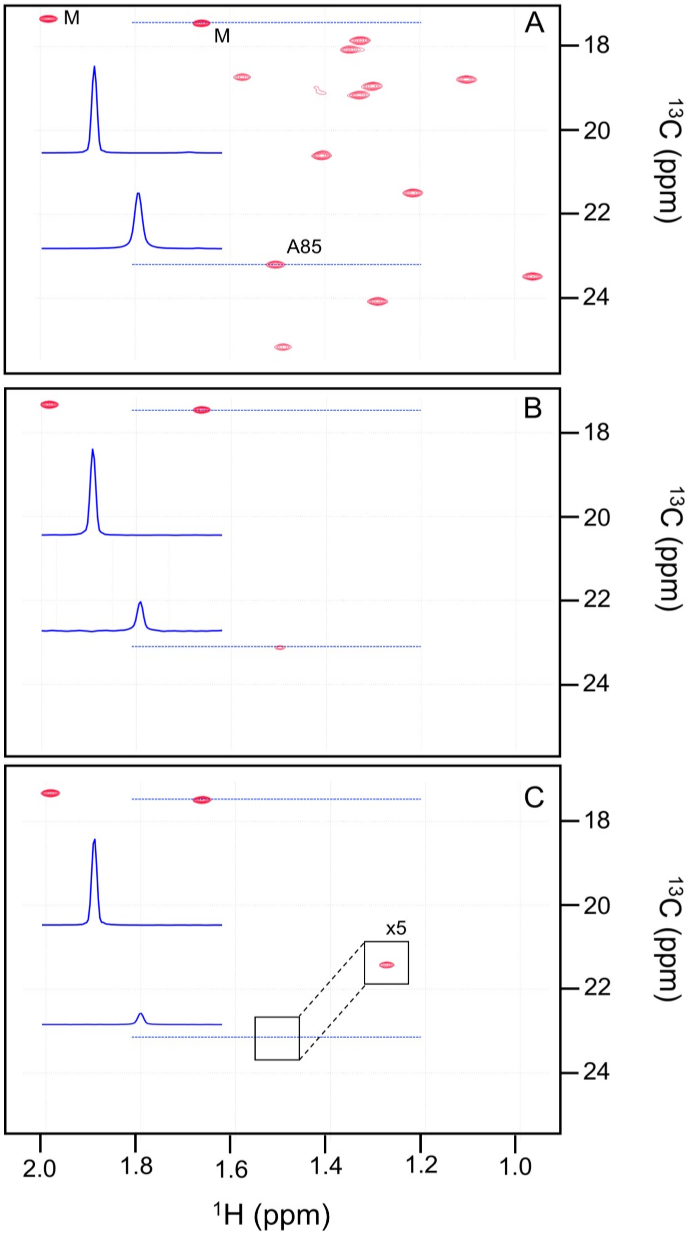


**Figure S2: 2D methyl-TROSY spectra of the HMA8 ATPase NBD** with **(A)** all methionine (2) and alanine (12) residues $^{13}CH_3$-labelled. For spectra presented in panel **(B)** and **(C)**, samples were $^{13}CH_3$-labeled on methionines and site-specifically labeled on A85 using G3:C70 mutated $tRNA_{CUA}$ **(B)** or wild-type $tRNA_{CUA}$ **(C)**. The first contours of all spectra are displayed at a level corresponding to 10% of the most intense methionine signal, except for the insert in panel **(C)**, which displays spectra at a 5-fold lower contour level. The 1D traces for a reference methionine signal and A85 are displayed in blue in each panel.

**Figure S3**

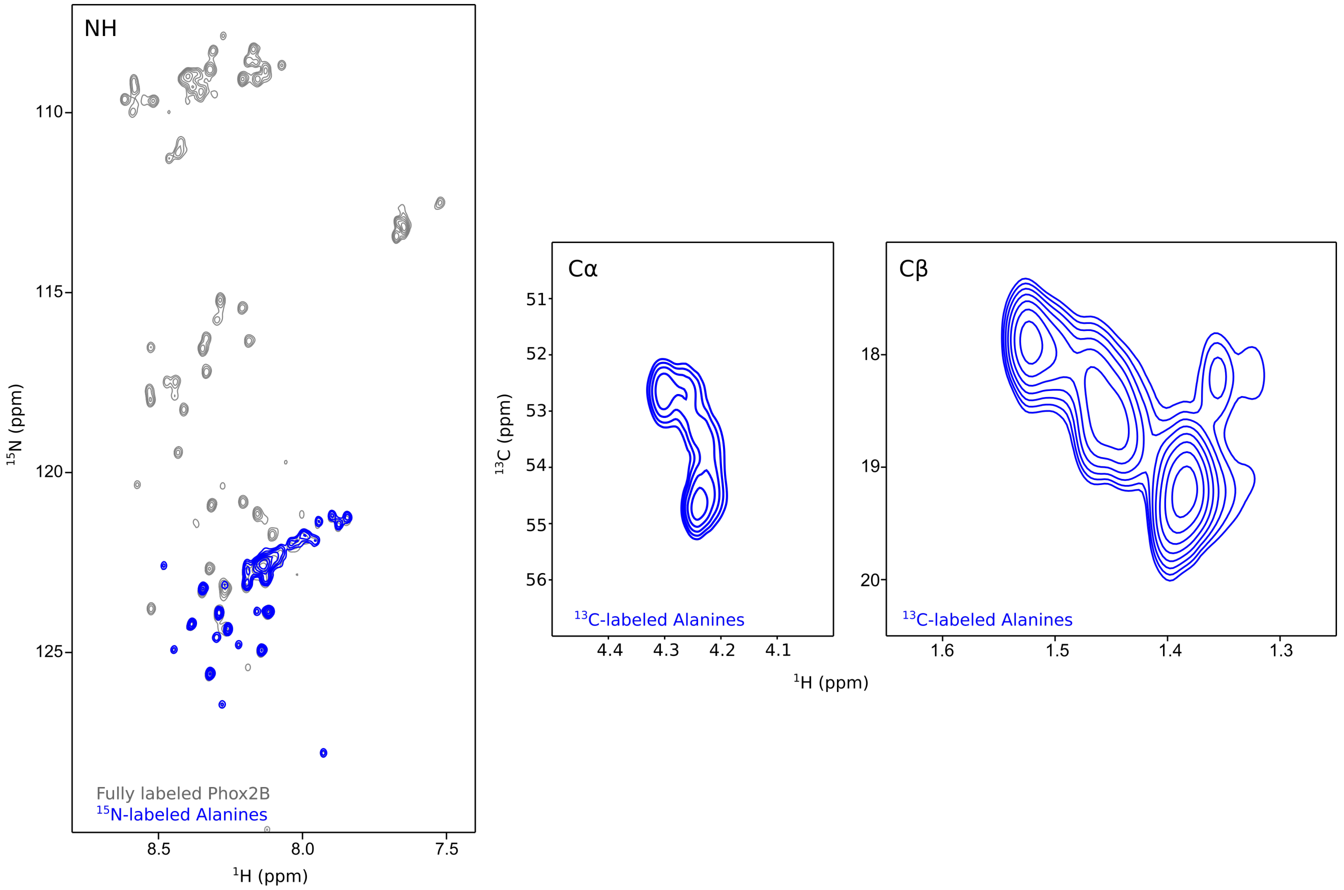


**Figure S3. NMR spectrum of Phox2B C-terminal domain.** (*Left*) $^{15}$N-HSQC spectrum of fully labeled Phox2B C-terminal domain (grey) overlayed to the only alanine-labeled one (blue). Cα (*middle*) and Cβ (*right*) alanine regions of the $^{13}$C-HSQC of the only alanine-labeled Phox2B C-terminal domain
